# Automatic Human-like Mining and Constructing Reliable Genetic Association Database with Deep Reinforcement Learning

**DOI:** 10.1101/434803

**Authors:** Haohan Wang, Xiang Liu, Yifeng Tao, Wenting Ye, Qiao Jin, William W. Cohen, Eric P. Xing

**Author notes:** Equal Contribution. The work is done while the author is at CMU.

## Abstract

The increasing amount of scientific literature in biological and biomedical science research has created a challenge in the continuous and reliable curation of the latest knowledge discovered, and automatic biomedical text-mining has been one of the answers to this challenge. In this paper, we aim to further improve the reliability of biomedical text-mining by training the system to directly simulate the human behaviors such as querying the PubMed, selecting articles from queried results, and reading selected articles for knowledge. We take advantage of the efficiency of biomedical text-mining, the flexibility of deep reinforcement learning, and the massive amount of knowledge collected in UMLS into an integrative artificial intelligent reader that can automatically identify the authentic articles and effectively acquire the knowledge conveyed in the articles. We construct a system, whose current primary task is to build the genetic association database between genes and complex traits of the human. Our contributions in this paper are three-fold: 1) We propose to improve the reliability of text-mining by building a system that can directly simulate the behavior of a researcher, and we develop corresponding methods, such as Bi-directional LSTM for text mining and Deep Q-Network for organizing behaviors. 2) We demonstrate the effectiveness of our system with an example in constructing a genetic association database. 3) We release our implementation as a generic framework for researchers in the community to conveniently construct other databases.

## 1. Introduction

Understanding the biological and biomedical science is one of the most fundamental goals of research and an essential step towards the realization of “precision medicine” in this era. Scientists all over the world are collaboratively contributing to this final goal, leading to an accompanying growth of the scientific literature. For example, PubMed^a^ has seen exponential growth regarding the number of publications in recent years^1^ and has collected over 27 million abstracts.^2^ These massive amount of articles consequently bring in the challenge of integrating the information conveyed effectively and accurately.

Biomedical information extraction has been the answer to this challenge for a long time.^3,4^ However, due to the demand of high reliability in biomedical research, following a typical general-purpose information extraction protocol and examining every article in the corpus nondiscriminatorily may lead to falsely constructed knowledge because of the non-negligible number of scientific literature with the issues of reproducibility.^5–7^

To fulfill the need of reliability in text mining and knowledge-base construction, instead of requiring the system to scan the entire corpus uniformly, we propose to train the system to directly simulate the behavior of a scientist with a sequence of actions including 1) querying the web, 2) evaluating the article, 3) studying the article for knowledge if necessary, 4) rejecting the knowledge if necessary, and 5) storing the knowledge. The 2^nd^ and 4^th^ steps play the essential roles in maintaining the reliability in constructed databases in our proposed system. Boosted by the power of deep reinforcement learning in organizing these actions, the ability of deep Bi-directional long short-term memory (LSTM) in text mining, and massive amount of knowledge encoded in Unified Medical Language System (UMLS),^8^ we are able to present our human-like system that can imitate the behaviors of a real scientist and construct the database of reliable and cutting-edge biomedical publications efficiently and endlessly. Therefore, we name our system the Everlasting Iatric Reader (Eir)^b^. We further apply our system to construct a genetic association database, where we can verify the performance of Eir with a manually crafted database of 167k gene-trait associations from high quality articles.^9^

The contributions of this paper are three-fold:

- We propose to improve the reliability of text-mining by building a system that can directly simulate the behavior of a researcher, and we develop corresponding methods, such as Bi-directional LSTM for text mining and Deep Q-Network for organizing behaviors.
- We demonstrate the effectiveness of our system with an example in constructing a genetic association database.
- We release our implementation as a generic framework for researchers in the community to conveniently construct other databases.

The remainder of this paper is organized as follows. In Section 2, we will introduce the related works in biomedical text mining. In Section 3, we will systematically introduce our system, mainly with deep reinforcement learning module that organizes the actions, text mining module that extracts the information, and implementation specifications. In Section 4, we will compare the performance to validate the strategy of Eir. Finally, in Section 5, we will draw conclusions and discuss about the future work.

## 2. Related Work

Text mining from biomedical literature has been studied extensively for a long time with a variety of different applications, such as patient analysis from electronic health records,^10–12^ gene annotations from protein networks,^13^ and drug repositioning from literature.^14^ One can refer to comprehensive reviews^4,15,16^ and the references therein for more detailed discussions.

The text mining usually leads to automatic construction of knowledge bases. In recent years, Mallory *et al.*^17^ curated a database of gene-gene interactions. They applied the information extraction engine DeepDive^18^ to around 100k full text PLOS articles for extracting direct and indirect gene-gene interactions. Poon *et al.*^19^ introduced the Literome project, where they extracted directed genic interactions and genotype-phenotype associations from PubMed articles. Lossio-Ventura *et al.*^20^ introduced a pipeline to build an obesity and cancer knowledge base. Very recently, Lossio-Ventura *et al.* also noticed the reliability issue of knowledge base, so they further proposed to incorporate cross-sourcing process to improve the reliability of the their previously developed knowledge base.^21^

On the other hand, the boom of deep learning techniques has allowed many more advanced methods developed for biomedical applications.^22–24^ As a result, LSTM and its variants,^25,26^ and word embedding techniques^27,28^ have been studied extensively for a variety of applications.

In comparison, a difference between most of previous work and our work is that we aim to improve the reliability of the extracted knowledge by examining the source unstructured data (*i.e.* the PubMed literature in our case). To put in simpler words, while most previous work are extending human’s intelligence of comprehending the articles, our system aims to extend human’s intelligence of the entire research process that starts with querying the web and selecting the interesting article. To the best of our knowledge, this paper is the first one that simulates the entire research process in biomedical information extraction to improve the reliability of the constructed knowledge base. However, many similar concepts^29–31^ have been proposed previously. Most relevantly, Kanani *et al*^32^ utilized reinforcement learning to reduce computational bottlenecks, minimizing the number of queries, document downloads and extraction action, a similar strategy has been proposed independently for biomedical text mining with the concept “focused machine reading”,^33^ which is inspired by Narasimhan *et al*,^34^ who built an information extraction system that can query the web for extra information with reinforcement learning.

## 3. Method

In this section, we officially introduce the our system. We will start with the main framework, and continue to introduce the deep reinforcement learning module that organize different actions of the system, which is followed by the discussions of proprecessing module and biomedical text mining module. After a systematic introduction of the detailed algorithms, this section is concluded with implementation specifications.

### 3.1 Model Framework

Eir’s research process is a markov decision process (MDP), where the model learns to query the search engine for scientific articles to read for the knowledge. We represent the MDP as a tuple < *S, A, T, R* >, where *S* = *s* is the space of all possible states, *A* = *a* is the set of all actions, *R*(*s, a*) is the reward function, and *T* (*s^′^|s, a*) is the transition function.

We present the details of these components as following:

- **Actions:** Action (we use *a* to denote action throughout this paper) is a set of Eir’s behaviors to simulate a real researcher, including

1. Query the search engine.
2. Evaluate whether the article is reliable.
3. Read the article for detailed information.
4. Exam credibility of the information and querying again.
5. Stop.

As shown in Figure 1, for every interesting query, Eir starts with the 1st action and then enters the loop from the 2^nd^ action to the 4^th^ action until Eir is satisfied with the finding of current research interest and ceases with the 5^th^ action. Then Eir repeats the entire process with another query.
- **States:** The state *s* in the MDP describes the research status of Eir, possible candidate states include the ones that are precedent or after each aforementioned action. There are only a countable number of actions, but we use continuous real-valued vector to represent each state so that we could have a better modeling power to distinguish Eir’s research status after each action. The state is constructed with a variety of information, including the embedding vector that the Bidirectional LSTM yields, the confidence of biomedical text mining module, the confidence of selecting an article to read, etc.
- **Rewards:**The reward function is chosen to maximize the intermediate paper selection accuracy and final extraction accuracy together while minimizing the number of queries. The accuracy component is calculated using the difference between the accuracy of the current and the previous set of entity values.
- **Transitions:**Transition *T* (*s^′^|s, a*) is modeled as a function of how the next state *s^t^* is updated given the current state *s′* and action *a* taken.

**Fig. 1:**
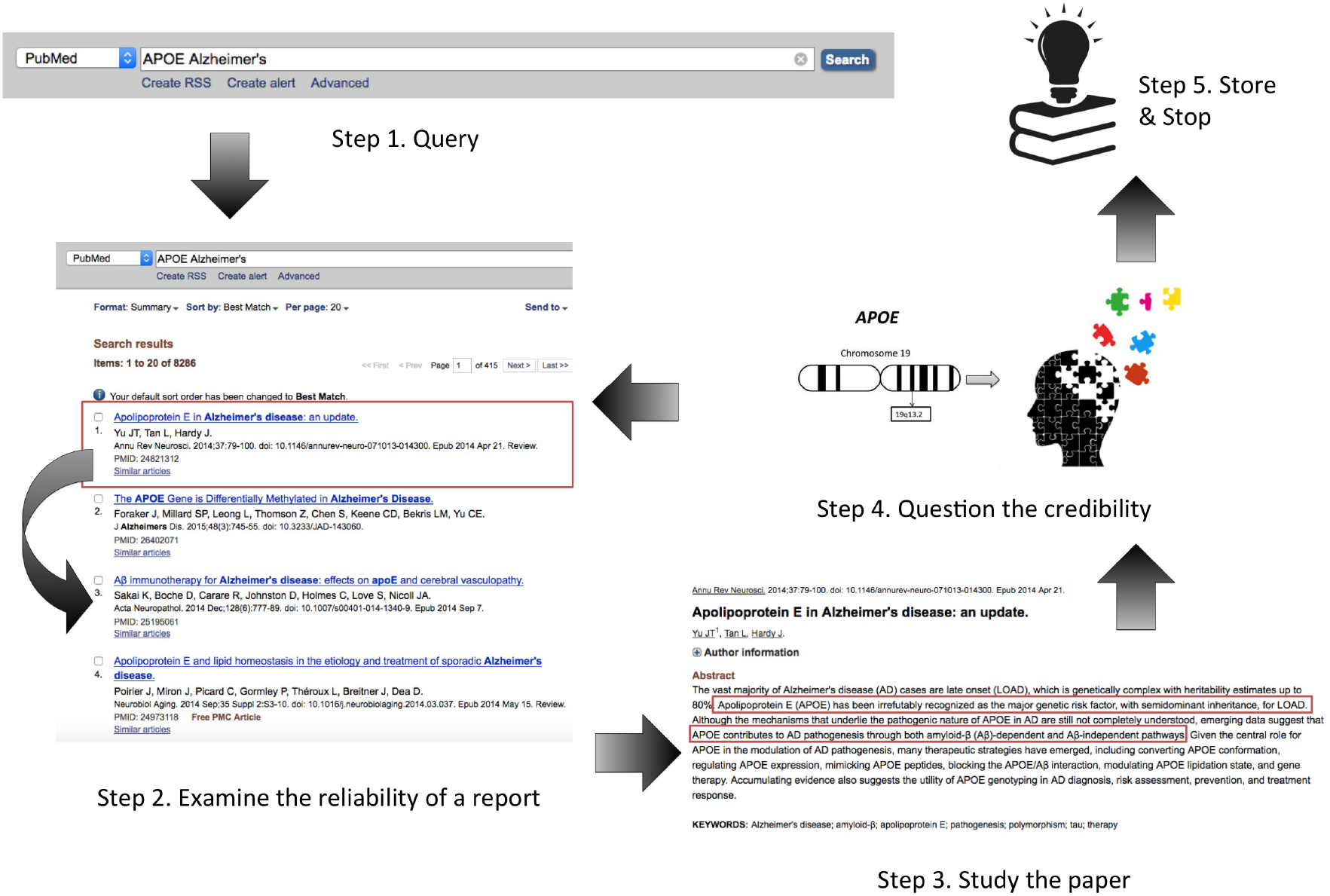
Overview of Eir’s possible behaviors

### 3.2. Deep Reinforcement Learning for Organizing Actions

As we have introduced previously, we utilized deep reinforcement learning to arrange the sequence of actions *a* to perform, given a state function denoted as *Q*(*s, a*). To update *Q*(*s, a*), we used the popular Q-learning,^35^ which iteratively updates *Q*(*s, a*) as following:

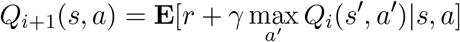

where *r* = *R*(*s, a*) is the reward and *γ* is a discounting factor.

Because of the continuous nature of our state space *S*, we use a deep Q-network (DQN)^36^ as a function approximator *Q*(*s, a*) = *Q*(*s, a*; *θ*). The Q-function of DQN is approximated by a neural network, whose parameters (*i.e. θ*) are updated through stochastic gradient descent. We followed the detailed parameter learning strategies introduced previously.^34^

### 3.3. Preprocessing and Name Entity Recognition with UMLS

Before we feed in the texts into the text mining module, we notice that the literature is filled with alternative, idiosyncratic and arbitrary names and symbols. The text mining module will only exhibit its full power when the texts are processed into a uniform representation. Therefore, we utilize the rich information collected by the unified medical language system (UMLS).^8^ UMLS defines a unique concept for all the terms that are interchangeable. For example, “Alzheimer’s disease”, “Alzheimer’s”, and “alzheimer” will be mapped into the same concept. UMLS contains over one million biomedical concepts that are split into 133 broad categories (such as “Organisms”, “Anatomical structures”, “Biologic function”). With the help of MetaMap,^37^ we are able to translate the unstructured texts into a sequence of concepts, together with the category information, an associated confidence score, and two binary values to indicate whether the concept is in gene ontology, and in disease ontology respectively.

### 3.4. Bidirectional LSTM for Relation Classification

As Eir queries PubMed with a gene-trait pair, the text mining model only needs to classify whether the returned texts from PubMed can be seen as evidences to support that there is association between the queried gene-trait pair. Therefore, the text mining module can be conveniently regarded as a classification module. We use a Bidirectional LSTM^38^ for classifying whether the text describes as association relationship between the gene and the trait the system queried. We choose this Bidirectional LSTM architecture mainly because we notice that it is empirically the best performing method among other neural architecture for our specific task. We first treat the sequence of concepts as words in text and created a 512-dimension vector of continuous values to represent each concept. Further, we feed in this concept-embedding, together with an one-hot representation of the category information, and the two binary values into the Bidirectional LSTM, which is trained through Adam.

### 3.5. Algorithm

Algorithm 1 describes the overall algorithm of the MDP process of Eir, where *g* and *t* stands for gene and trait respectively, *a* stands for action, *s* stands for state, and *r* stands for reward. “Agent” refers to DQN, which organizes the sequence of actions given states and reward. Details including the methodology of updating (*s, r*) has been discussed in previous sections.

### 3.6. Implementation Specification

The Deep Reinforcement Learning component of Eir is implemented as an extension of Narasimhan *et al*,^34^ we also use a DQN consisting of two linear layers (20 hidden units each) followed by rectified linear units (ReLU), along with two separate output layers.

The web query component is built with a web crawling engine Scrapy^c^ communicating with NCBI PubMed search engine. At this moment, we only query for the abstracts of the articles. We only work with abstracts for three reasons: 1) this allow us to conveniently access and scan a large amount literature, 2) we notice that a majority of articles disclose the most important findings in the abstract with a straightforward style of writing, 3) previous work notice that mining from full texts may lead to more false positives.^39^

The preprocessing module is built as a python script that runs MetaMap, which is a binary software that allows users to conveiently annotate words and phrases of texts with manually defined concepts in UMLS.

The sentences are truncated with max length of 300 concepts. We only consider the 30,000 most frequent concepts together with the specific defined ‘SOS’ (start of sentence), ‘EOS’ (end of sentence), ‘UNK’ (unknown) and ‘PAD’ (padding the sentences shorter than 300) concepts. We use a 2-layer Bidirectional LSTM with hidden dimension set to 1000, and feed 512 dimension concept embedding, one dimension gene ontology, one dimension disease ontology, and 136 dimension semantic type as the input of LSTM. The LSTM is trained jointly with the embedding matrix using Adam with step size set to 0.00004 and batch size set to 64.

Then We train the Eir models for 10000 steps every epoch using the Maxent classifier as the base extractor, and evaluate on the entire test set every epoch. The final accuracy reported are averaged over three independent runs; each runs score is averaged over 5 epochs after 45 epochs of training. The penalty per step is set to −0.001. We used a replay memory of size 500k, and a discount *γ* of 0.8. We set the learning rate to 2.5E5. The *ϵ* in *ϵ*-greedy exploration is annealed from 1 to 0.1 over 500k transitions. The target-Q network is updated every 5k steps. The whole framework was trained to optimize the reward function.

We release our implementation^d^ for the community to use our system or build more advanced text mining module into our system for better performance.

**Algorithm 1.**
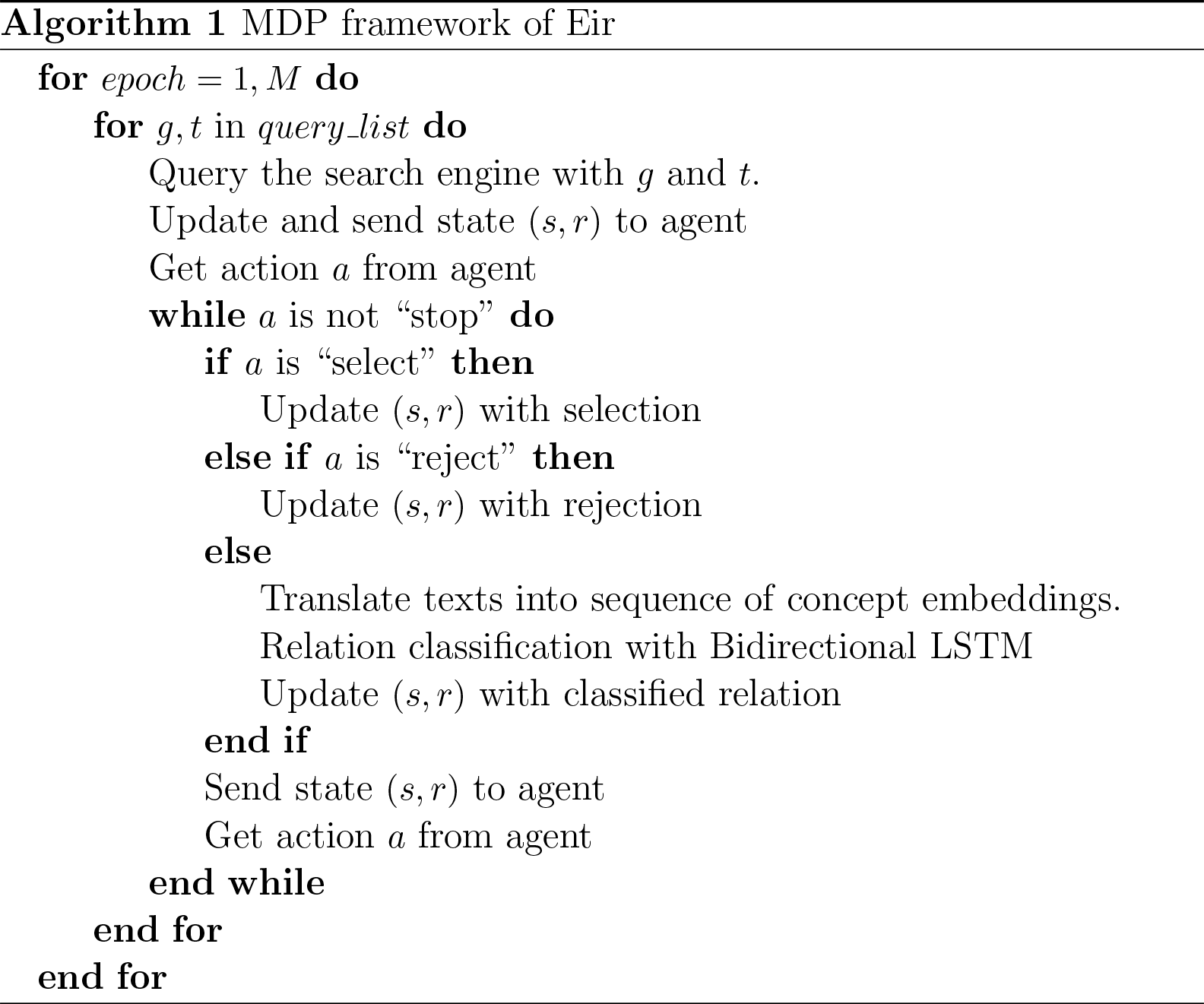
MDP framework of Eir.

## 4. Experiments

In this section, we will verify the performance of Eir by showing that, with the same text mining module, the Eir system can help improve the performance of extracted associations. We will first discuss how we construct the experimental data sets then discuss the results.

### 4.1. Data

Within the scope of this paper, Eir focus on constructing the knowledge base for gene-trait association relationship of human. To enable Eir to learn the associations, we utilized the high quality data set of 167k association relationship that is manually crafted for over ten years.^9^

In addition to the gold-standard information of gene-trait association relationship, another contribution of this data set is the collection of high quality publications that report these associations. Every entry in the database is grounded by the authentic source of scientific paper that originally publishes the relationship. These detailed information grants us the possibility of directly training Eir to discriminate the reliable papers out of the less favourable papers that were not selected by GAD curators for some reasons.

Despite that Eir is designed for extracting latest information online, in order to test the effectiveness of Eir, we need to run the core functions on a local collections of articles with manually labelled true associations. Therefore, we query the PubMed with 54,041 queries of gene-trait pairs through our API and download 913,939 results with 305,651 distinct medical articles. After removing some invalid records (e.g. articles with invalid PMID), there are roughly 133,548 records (44,592 distinct articles) appear in the GAD database, which will serve as the reliable articles. As the construction of GAD ceased in 2014, we regard the articles that are published before 2014 but not in the GAD database as less favorable articles. To balance the data set for performance evaluation, we sampled 140,361 less favorable records before 2014 for comparison. Note that, these less favorable articles are not collected randomly, but are returned from PubMed search engine when we query with a pair of gene and trait. Besides, we delete the articles whose titles and abstracts do not contain the queried gene and trait explicitly to remove obviously irrelevant articles. Then, we random split the whole data set to sample 80% records as training data, and the rest as testing data. The training set consists of 55k records, the testing set consists of 219k records.

### 4.2. Evaluation

In order to show the effectiveness of the Eir system, we compare the system’s precision, recall, and F1 score with a conventional biomedical text mining strategy that scans all the documents nondiscriminatorily. As Eir uses the Bidirectional LSTM for text mining module, we use the same model as baseline method for fair comparison.

### 4.3. Results

#### 4.3.1. Improved Reliability

We first train our baseline Bidirectional LSTM and the results are shown in the Table 1 (first row). The Bidirectional LSTM yields a precision of 91.25%, a recall of 96.55%, and an overall F1 of 93.80%. These numbers indicate that the Bidirectional LSTM is capable to capture the feature of authentic articles.

**Table 1:** Results of Reliability Comparison

|  | precision | recall | F1 |
| --- | --- | --- | --- |
| Bidirectional LSTM | 91.25% | 96.55% | 93.80% |
| Eir | 91.4% | 97.0% | 94.1% |

Further we add the Deep Reinforcement Learning component to train the overall Eir system. The results of Eir are shown as Table 1 (second row). We can see that the precision score is 91.4%, the recall score is 97.0%, the F1 score is 94.1%. Compared to the baseline model, our Eir framework is better at extracting the features of valuable articles and utilizing the information and can retrieve the authentic articles more efficiently by employing the Deep Reinforcement Learning module.

#### 4.3.2. Robustness in Real-world Situations

To better simulate the real-world situation that the researchers are in, we remove different percentage of authentic articles both in the training data set and in the testing data set, for the researchers get ample amount of less favorable articles. We randomly remove a certain percentage of authentic articles to do the ablation experiments. As the percentage of authentic articles decreases, the difficulty of our task increases. The results are shown in Table 2. We can see the Eir system is more robust than the baseline model under these situations. Eir reports higher precision, recall, and F1 score in all of these settings. More interestingly, we calculate the increments Eir achieves over baseline model. We notice that, as the difficulty increases, the increment also increases. Therefore, we believe Eir will be more helpful in the real-world situation when a large amount of articles are less favorable articles.

**Table 2:** Results of Eir in real-world situations

|  | Full Data |  |  | 20% Authentic Articles |  |  | 10% Authentic Articles |  |  |
| --- | --- | --- | --- | --- | --- | --- | --- | --- | --- |
|  | Prec | Recall | F1 | Prec | Recall | F1 | Prec | Recall | F1 |
| Bi-LSTM | 91.25% | 96.55% | 93.80% | 87.7% | 95.7% | 91.5% | 86.9% | 92.2% | 89.4% |
| Eir | 91.4% | 97.0% | 94.1% | 87.9% | 96.9% | 92.2% | 87.8% | 96.9% | 92.1% |
| Increment | 0.16% | 0.47% | 0.32% | 0.23% | 1.25% | 0.77% | 1.04% | 5.10% | 3.02% |

#### 4.3.3. Number of Articles Read

Finally, we examine Eir’s performance in the numbers of articles it needs to read to make a decision. Since Eir stops once it believes it has sufficient amount of information, we anticipate Eir will inspect less amount of articles than baseline models. To conduct this experiment, we exclude the gene-trait query pair with only one authentic articles. In the remaining data set, there is on average 2.54 articles for every query, and Eir reads only on average 2.46 articles. We further repeat this experiment with a data set that excludes all the articles with less than 4 articles per query, resulting in a data set with on average 6.23 articles for every query. Eir reads on average 6.10 articles.

## 5. Conclusions and Future Work

In this work, we introduced a system, namely Everlasting Iatric Reader (Eir), for biomedical text mining. A distinct difference between our system and previous biomedical text mining works is that our system is aimed to directly simulate the behaviors of scientists, including searching for scientific literature, examining the reliability of the manuscript, studying the paper for details, and continuing to search with suspicion of the learned knowledge.

In contrast to traditional biomedical text mining tools, the distinguishable advantage Eir has is the ability to discriminate reliable articles out of questionable articles and to shield the problems introduced by humans. This ability is particularly important in biomedical areas because in clinics, a falsely constructed knowledge may lead to fatal errors, while a missing piece of true knowledge will at most delay the cure of certain disease. Also, it is necessary to select trustworthy papers to read for information because it is known that there is a non-negligible number of publications with the troubles of reproducibility.

There are also limitations of the current Eir. For example, the action of Eir for evaluating the literature quality is trained supervisedly. The performance of our Eir can be greatly improved with a more cleaned data source, as now the false positives are introduced by some manuaaly crafted data that are labeled not correctly. Therefore, we will need a manually crafted data set first before we use Eir in some application. In this paper, we choose to construct the genetic association database because of the availability of GAD.^9^ However, there are still a large number of manually curated databases with information about which paper these information comes from, such as GWAS Catalog^40^ for SNP-phenotype association or UniProt^41^ for protein function annotation.

Looking into the future, a direct extension of our work is to broaden Eir vision to ask investigate into more biomedical topics in addition to gene-trait association relationships. Our immediate next-step plan is to train Eir for SNP-phenotype association with GWAS Catalog, then we can integrate these databases into GenAMap,^42^ a visual machine learning tool for GWAS^ [^e^ http://www.genamap.org/]^, for validation purpose of GWAS results. On the method development side, we hope to upgrade the biomedical text mining module with state-of-the-art methods to improve the information extraction performance, so that Eir could serve the community better. As a long-term plan, we hope Eir could help the community to build the omini-biomedical knowledge base, therefore, we released the source code of Eir for others in the community to use.

## 6. Acknowledgement

This work is funded and supported by the Department of Defense under Contract No. FA8721-05-C-0003 with Carnegie Mellon University for the operation of the Software Engineering Institute, a federally funded research and development center. This work is also supported by the National Institutes of Health grants R01-GM093156 and P30-DA035778.

## Footnotes

a the database maintained by the National Center for Biotechnology Information (NCBI)

b We name our system Everlasting Iatric Reader because it can endlessly construct the knowledge in the medical area, where the high reliability is an issue, and also because the acronym (Eir) shares the name of the goddess of medical knowledge in Norse mythology, which is related to the final goal of this and following-up projects.

c https://scrapy.org/

d https://github.com/lebronlambert/Eir

